# Local adaptation to an altitudinal gradient: the interplay between mean phenotypic trait variation and phenotypic plasticity in *Mimulus laciniatus*

**DOI:** 10.1101/2023.08.02.551729

**Authors:** Jill M. Syrotchen, Kathleen G. Ferris

**Affiliations:** Department of Ecology and Evolutionary Biology, Tulane University, 6823 St. Charles Avenue, New Orleans, LA 70118

**Keywords:** phenotypic plasticity, local adaptation, heterogeneity, elevation gradient, trait variation, leaf lobing

## Abstract

Organisms can adapt to environmental heterogeneity through two mechanisms: (1) expression of population genetic variation or (2) phenotypic plasticity. In this study we investigated whether patterns of variation in both trait means and phenotypic plasticity along elevational and latitudinal clines in a North American endemic plant, *Mimulus laciniatus*, were consistent with local adaptation. We grew inbred lines of *M. laciniatus* from across the species’ range in two common gardens varying in day length to measure mean and plastic trait expression in several traits previously shown to be involved in adaptation to *M. laciniatus’s* rocky outcrop microhabitat: flowering time, size-related traits, and leaf shape. We examined correlations between the mean phenotype and phenotypic plasticity, and tested for a relationship between trait variation and population elevation and latitude. We did not find a strong correlation between mean and plastic trait expression at the individual genotype level suggesting that they operate under independent genetic controls. We identified multiple traits that show patterns consistent with local adaptation to elevation: critical photoperiod, flowering time, flower size, mean leaf lobing, and leaf lobing plasticity. These trends occur along multiple geographically independent altitudinal clines indicating that selection is a more likely cause of this pattern than gene flow among nearby populations with similar trait values. We also found that population variation in mean leaf lobing is associated with latitude. Our results indicate that both having more highly lobed leaves and greater leaf shape plasticity may be adaptive at high elevation within *M. laciniatus.* Our data strongly suggest that traits known to be under divergent selection between *M. laciniatus* and close relative *Mimulus guttatus* are also under locally varying selection within *M. laciniatus*.

## INTRODUCTION

Environmental heterogeneity can come in multiple forms, such as spatial and temporal variation in resource availability (Cortés and Wheeler 2018). This heterogeneity is a driver for numerous ecological and evolutionary processes, such as community composition, species richness, and local adaptation (Stein, Gerstner, and Kreft 2014; Yang et al. 2015; Forester et al. 2016). Where environments vary, populations with enough genetic variation have the opportunity to adapt to their local conditions via natural selection and this is a common phenomenon across many taxa (Via and Conner 1995; Klich 2000; Hereford 2009; Nunez et al. 2021). Local adaptation occurs when an organism’s trait expression optimizes fitness values for its local environment, and is achieved by low gene flow and strong selection between populations (Kawecki and Ebert 2004). Mean phenotypes, here defined as the average expression of a trait due to genetic differences in a common environment, have been the primary focus of studies in the local adaptation literature (Hereford 2009; Savolainen, Lascoux, and Merilä 2013). Phenotypic plasticity, or the ability of one genotype to express multiple phenotypes in response to the environment, has also been hypothesized to be locally adaptive, particularly in response to predictable environmental heterogeneity (Snell-Rood and Ehlman 2021). In order for phenotypic plasticity to respond to locally varying natural selection there must be genetic variation in plasticity, or a genotype by environment interaction (GxE) (Josephs 2018). Therefore, demonstrating that a plastic trait exhibits GxE is the first step in investigating whether phenotypic plasticity is locally adaptive. Although classic studies in *Impatiens campensis* have detected an adaptive role for plasticity (Donohue et al. 2000, 2001), this has been difficult to test in most systems (Thompson 1991; Via et al. 1995; Donohue et al. 2001; Scheiner 2013).

In a natural landscape, mean phenotypic expression and phenotypic plasticity both contribute to an individual’s lifetime fitness, and therefore both have the potential to play a role in local adaptation. It can be difficult to identify when plasticity is locally adaptive in natural populations because the amount of plasticity in a trait must be measured using the same genotype in different environmental conditions. These constraints can be overcome with experimental designs using clonal replicates, or half-siblings in systems where inbreeding or cloning cannot be performed (Relyea 2002; Cooper et al. 2022). Plants are sessile and many of them can be inbred or replicated clonally making them ideal for investigating the relative contributions of mean phenotype and plastic trait expression to local adaptation.

Characterizing the plastic response of a species to environmentally relevant conditions is important in understanding its life history (Wilczek et al. 2009). Sarkar (2004) postulated that, if tested for, plasticity is bound to be present in different environments and yet there are still unanswered questions about the variation in, mechanisms of, and significance of phenotypic plasticity in most natural populations. While plasticity in a single trait has been examined in numerous systems (Chevin and Hoffmann 2017), few studies have examined plasticity in multiple traits across an organism’s lifetime. Therefore we lack data about the degree of correlation between plastic expression of multiple traits and how these correlations may influence fitness (Nielsen and Papaj 2022). Although the genetic basis of phenotypic plasticity has been mapped in traits like flowering time (Fournier-Level et al. 2022) and branching pattern in *Arabidopsis* (de Jong et al. 2019), whether plastic variation in a given trait is controlled by separate genetic loci from variation in the trait mean is still not well understood (but see Fournier-Level et al. 2022). By comparing the correlation between plastic and mean phenotypic trait expression in a population exhibiting GxE we can investigate whether plasticity has independent genetic controls (Holeski, Chase-Alone, and Kelly 2010). This information is foundational in determining how plasticity is expressed from genotype to phenotype.

Elevation and latitude exert spatially varying selection across populations. Due to their sessile nature, plants experience strong natural selection based on the environment around them. Local adaptation to elevation has been found in many plant systems such as in *Arabidopsis arenosa* populations from four foothill-alpine clines (Wos et al. 2022) and in *M. guttatus* in response to drought stress (Kooyers et al. 2015). Phenotypic plasticity is known to be involved in local adaptation in *Populus fremontii* populations across Arizona, where plasticity in traits such as bud set and plant height were found to vary along an elevational gradient (Cooper et al. 2022). Local adaptation of phenotypic plasticity in a number of plant functional traits in *Phragmites australis* was found to follow a latitudinal gradient (Ren et al. (2020).

One important environmental cue in plants that may drive local adaptation is photoperiod. Photoperiod varies temporally both predictably and unpredictably during the growing season for sessile alpine and subalpine organisms. Photoperiod varies predictably with shorter days occurring in the winter and the longest days occurring at the summer solstice. In this way, photoperiod is a reliable sensing cue, which is hypothesized to be important for the development of phenotypic plasticity (Snell-Rood and Ehlman 2021). Elevation and latitude are two environmental gradients that interact with photoperiod in a heterogeneous manner as a result of annually varying temperatures and snowpack. High latitude populations will always experience longer photoperiods during the spring and summer growing seasons than low latitude populations. Experienced photoperiod varies less predictably in mountainous environments (Figure 1a and 1b). High elevations accumulate more snowfall and have a later timing of snowmelt than low elevations, which means that for plants growing at the same latitude low elevation populations will germinate and develop under shorter photoperiods than those occurring at higher altitudes. However over time the amount of snowpack varies significantly meaning that the experienced photoperiod of a short-lived annual plant can fluctuate from year to year in alpine environments (Bormann et al. 2018). Therefore, photoperiod may be an important cue that young seedlings utilize in alpine environments to determine time of year and their subsequent developmental trajectory.

**Figure 1.**
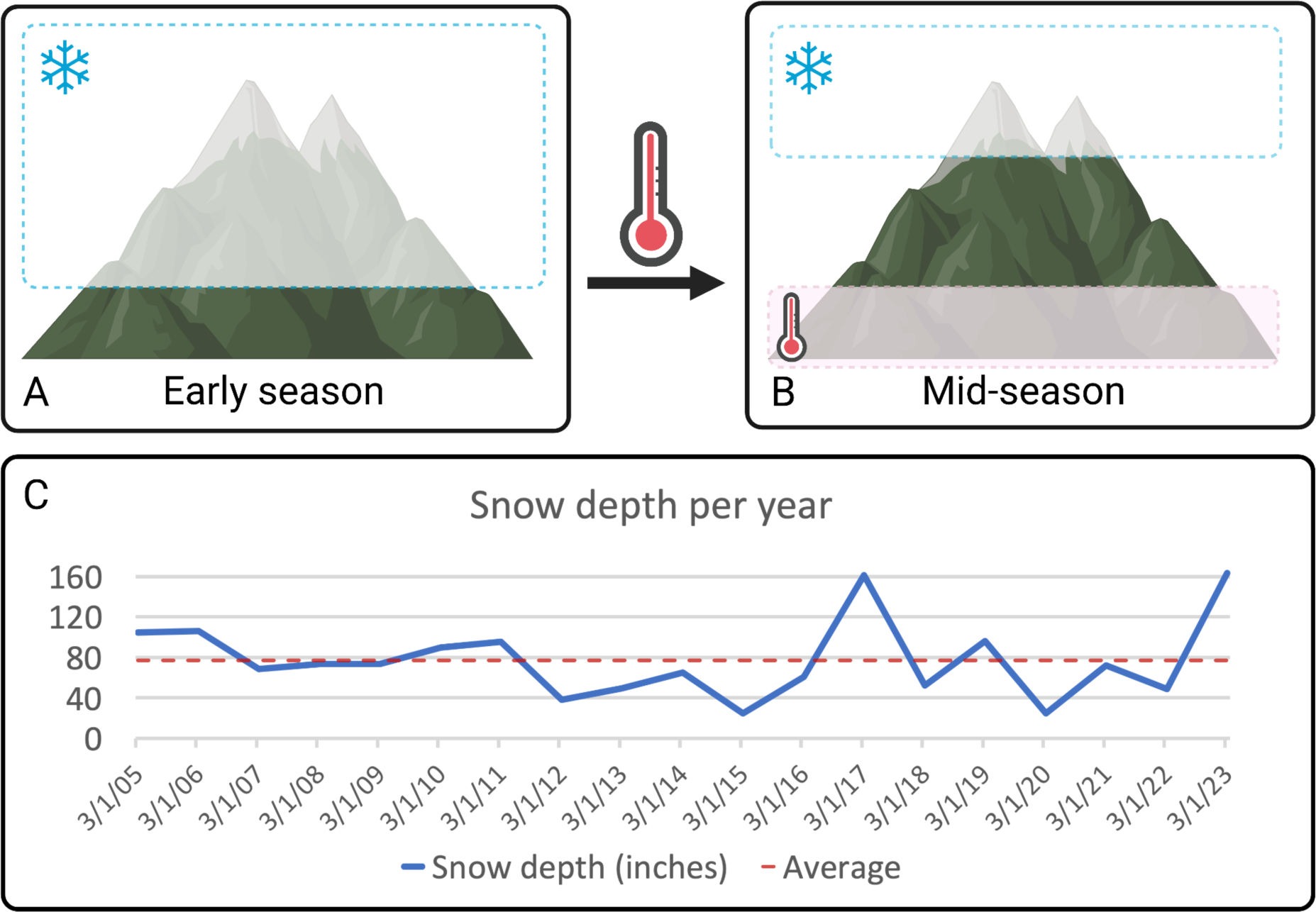
a) Early in the growing season, only lower elevation populations have warmed up enough to experience complete snowpack melt. These populations begin to grow earlier in the spring, and therefore experience a shorter photoperiod. b) By early to mid summer, all snowpack has melted from higher elevation sites, therefore high elevation populations grow during longer photoperiods. c) Snowpack level changes from year to year, and the pace of warming may be slower or faster. This creates fluctuation in experienced photoperiod over time as well as across elevation. Snow depth data collected at Dana Meadows in Yosemite National Park (California Data Exchange Center 2023). Created with BioRender.com.

*Mimulus laciniatus*, a member of the *M. guttatus* species complex, is a Sierra Nevadan endemic plant that grows on rocky granite outcrops from 1000-3300 meters in elevation (Fenster and Ritland 1994; DeMarche, Rice, and Sexton 2013). Depending upon elevation and the timing of snow melt, *M. laciniatus* populations experience different day lengths during the growing season, making this an interesting system to examine patterns of local adaptation in response to photoperiod fluctuation. A previous study by Ferris and Willis (2018) found that flowering time, plant height, and leaf shape were under divergent selection between *M. laciniatus* and its close relative *M. guttatus*. Leaf shape, and in particular leaf dissection, has been found to vary clinally across elevation and latitude within other plant species (Gurevitch 1988; Campitelli and Stinchcombe 2013; Wong et al. 2020). In many species it is unclear whether this trait variation and potential adaptation arises via genetic variation in mean phenotype expression, phenotypic plasticity, or a combination of the two (but see Bright and Rausher (2008); Campitelli and Stinchcombe (2013)).

In this study we investigated whether mean and plastic variation in leaf shape, flowering time, and floral size among *M. laciniatus* populations are consistent with local adaptation to elevation and latitude. We grew replicates of genotypes from twenty-one populations across *M. laciniatus’* elevational and latitudinal range in long and short day common garden experiments. We then tested whether patterns of mean and plastic trait expression were associated with elevation or latitude in the population of origin. This allowed us to examine whether the same traits that are under divergent selection between closely related sympatric species may also be under locally varying selection within *M. laciniatus*.

## MATERIALS AND METHODS

### Study system

*Mimulus laciniatus* is a small, self-fertilizing annual plant, with low levels of gene flow expected between most populations (Ferris, Sexton, and Willis 2014; Sexton et al. 2016). Unlike *M. guttatus* which displays rounded leaves, *M. laciniatus* has deeply lobed, narrow (laciniate) leaves as a defining characteristic of the species (Ferris et al. 2015). We created inbred lines from genotypes collected from natural populations of *M. laciniatus* across its geographic range to investigate phenotypic responses to photoperiod fluctuation. Due to its high rate of self-fertilization (93-97%) *M. laciniatus* is already highly inbred when first collected from the field (Ferris et al 2014). We self-fertilized each genotype for at least 5 additional generations in the lab. Twenty-one populations of *M. laciniatus* from across the central and southern Sierra Nevada mountain range (California, USA) were used in this experiment (Table 1, Figure 2). These populations originate from across 1000-2500m in elevation and two degrees of latitude (∼36°-38°; Figure 2) and six distinct watersheds (Figure S1). Low levels of gene flow between populations have been observed in previous studies of *M. laciniatus* (Ferris, Sexton, and Willis 2014; Sexton et al. 2016).

**Figure 2.**
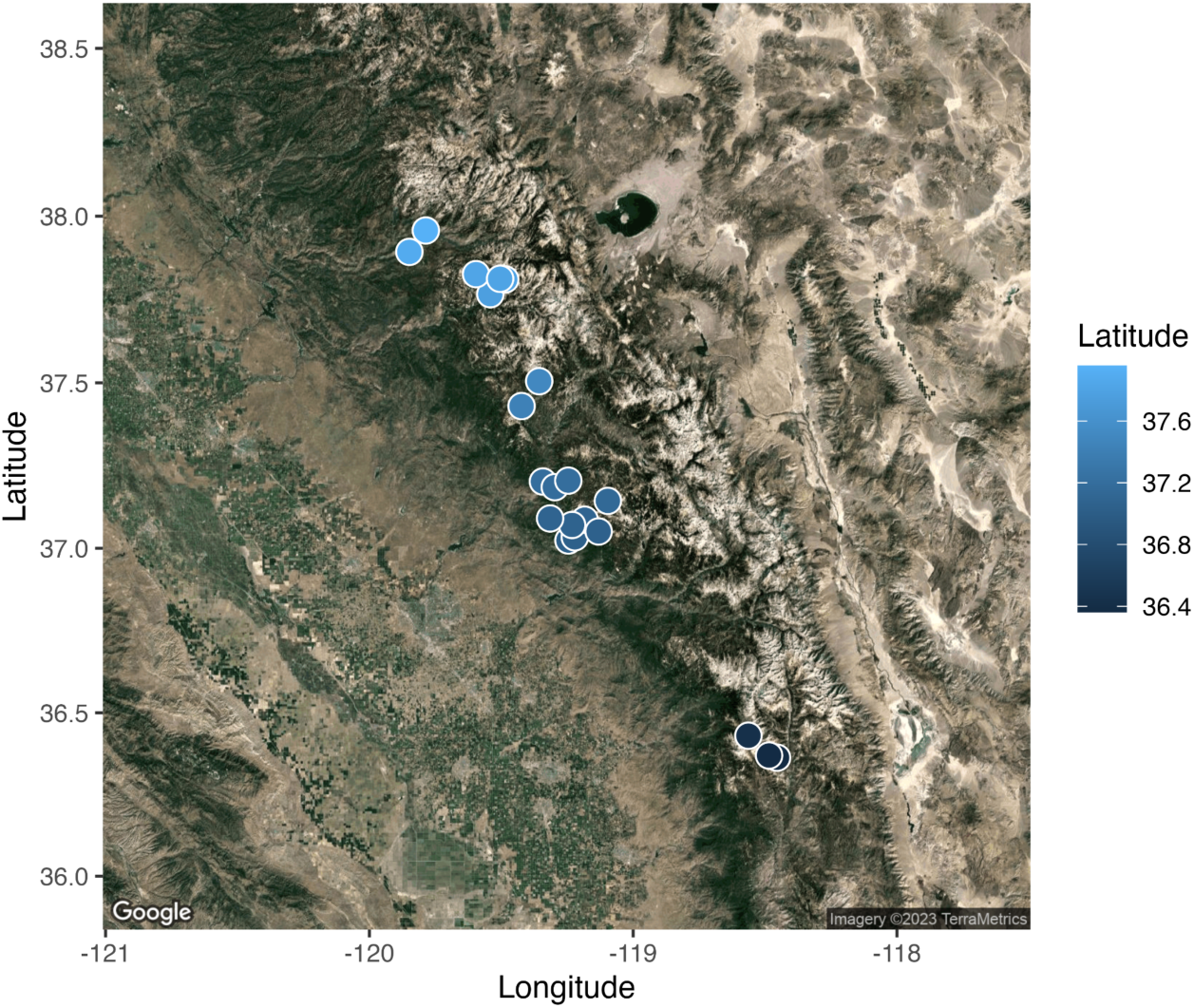
A map of the Sierra Nevada mountain range with each population mapped to its original collection location. Populations are colored based on latitude.

**Table 1.** Geographic data for each *M. laciniatus* population used in the study.

| Population name | GPS coordinates (Decimal Degrees) | Location description | Elevation (m) |
| --- | --- | --- | --- |
| BCL | 37.2049, -119.246 | Big Creek, Fresno County, CA | 1493.45 |
| BML | 37.090509, -119.3150 | Mill Run Lane, Shaver Lake, CA | 1696.74 |
| CECL | 37.300143, -119.4835 | Central Camp Rd, Madera County, CA | 1274.67 |
| CRV | 37.812585, -119.50447 | South of Olmstead Point parking area, Yosemite National Park, CA | 2577.44 |
| DNK | 37.0511, -119.13102 | Dinky Creek Rd, Sierra National Forest, CA | 1851.57 |
| GB | 37.0708, -119.23165 | Grand Bluff off Hwy 168, Sierra National Forest, CA | 1679.37 |
| HEH | 37.89314, -119.84853 | Evergreen Rd, Yosemite National Park, CA | 1396.83 |
| HHR | 37.957206, -119.7860 | Hetch Hetchy Reservoir trail, Yosemite National Park, CA | 1207.56 |
| HUL | 37.14372, -119.09606 | Hwy 168 before south turn off to Huntington Lake Rd, Fresno County, CA | 2170.68 |
| LIB | 36.36958, -118.48498 | Generals Hwy east of Little Baldy, Sequoia National Park, CA | 2196.28 |
| OPN | 37.8107, -119.4852 | Olmstead Point, Yosemite National Park, CA | 2529.72 |
| PETL | 37.03346, -119.22122 | Peterson Rd, Fresno County, CA | 1256.63 |
| QAL | 36.7203, -118.02306 | By Quail Flat off Generals Hwy, Tulare County, CA | 1986.99 |
| SBL | 37.02384, -119.2455 | Sandy Bluffs, Hwy 168, Sierra National Forest, CA | 991.77 |
| SHL | 37.08682, -119.18388 | Hwy 168, Shaver Lake, CA | 1594.33 |
| ST | 37.76637, -119.54167 | Snow Trail near falls, Yosemite National Park, CA | 1869.95 |
| TRT | 37.42990, -119.42321 | Hwy 41 below Turtleback Dome, Yosemite National Park, CA | 1480.65 |
| WCL | 37.20227, -119.34225 | Willow Creek by Bass Lake Rd, Fresno County, CA | 1100.27 |
| WLF | 37.50492, -119.35631 | Tioga Rd east of White Wolf, Yosemite National Park, CA | 2395.31 |
| WUV | 36.681, -118.88028 | Along road to Wuksachi Village, Sequoia National Park, CA | 2129.638 |
| YCN | 37.8264, -119.5957 | Yosemite Creek on Tioga Rd, Yosemite National Park, CA | 2346.85 |

### Experimental design

To compare the variation in mean phenotype and plastic expression of quantitative traits among *M. laciniatus* populations from across the species range we grew genetic replicates in 1) a common garden in a long day environment to measure mean phenotypic expression of traits at UC Davis in 2017, 2) another long day common garden experiment at Tulane University in 2020, and 3) a common garden in short days at Tulane University in 2021 to compare to the Tulane long day garden and measure plasticity. We repeated the long day garden at Tulane to control for possible effects of chamber and location on our phenotypes when measuring plasticity. By replicating our long day garden, we are also able to avoid the statistical artifact of regression to the mean when testing for differences in patterns of variation in mean phenotype vs. phenotypic plasticity (Gunderson and Revell 2022). To measure genetic variation in mean expression in a single environment we grew plants at UC Davis in a Conviron walk-in chamber under 15 hour days at 21°C. To measure phenotypic plasticity, we sequentially grew plants at two different photoperiods (11 hour & 15 hour days) in a Conviron walk-in chamber at Tulane University. These long (15 hour) and short (11 hour) photoperiods represent the longest and shortest natural day lengths within *M. laciniatus’* native range.

We grew three to ten unique inbred lines from each of twenty-one populations for a total of 120 inbred lines in each growth chamber experiment. Seventy-one of those inbred lines were grown in both the mean phenotype experiment at UC Davis and plasticity grow-outs at Tulane. Three to five replicates of each inbred line per grow-out gave us a total of 1015 plants used to measure mean expression of traits and 677 plants used to measure plasticity in this study. We planted seeds atop 3” pots filled with wetted Sun Gro Fafard 3B/Metro-Mix 830 soil. Pots were stratified in dark Percival chambers set at 4°C for 10 days and misted every other day. After 10 days, we moved pots to a Conviron walk-in growth chamber set to 21°C. We kept pots constantly wet on full flood benches and misted daily. After seedlings showed their first set of true leaves, we flooded benches with water for 8 hours (during light hours) and kept them empty for 16 hours daily.

### Phenotypic measurements

To examine ecologically relevant phenotypic variation in *M. laciniatus* across populations, inbred lines, and photoperiods, we measured traits that have previously been shown to be under selection in *M. laciniatus’* native habitat: flowering time (days from germination to first flower), leaf lobing, and plant height, as well as mating system-related traits. The specific mating system traits measured were: corolla length, corolla width, longest stamen length, shortest stamen length, pistil length, and pseudo-cleistogamy (denoted as either pseudo-cleistogamous for a closed flower or chasmogamous for an open flower). *Mimulus laciniatu*s is pseudo-cleistogamous rather than truly cleistogamous because while the size and shape of the outer corolla changes, the internal structure of the plant does not (Lord 1981). Table S1 provides a chart of all traits measured. Critical photoperiod was also measured and calculated as the proportion of individuals per genotype that reached flowering under the 11h photoperiod. All plants flowered under 15h. For plants that flowered, we collected all phenotypic data on the day of first flower. For plants that did not flower under 11 hours, but instead maintained a vegetative state, we collected leaf lobing and plant height data two months after germination. We collected leaf lobing data by harvesting the most lobed leaf from the second node of each plant. We taped leaves to a sheet of white paper and scanned them for digital convex hull analysis of leaf lobing in ImageJ (Ferris et al. 2015). The convex hull is the shape that is created by connecting the outermost points of a leaf. The lobing index provided by convex hull analysis is calculated by dividing the difference between the area of the true leaf and the area of the convex hull by the larger area of the convex hull (Figure S2). The lobing index is a metric of leaf lobing from 0 to 1, with a higher number equaling higher lobing (Ferris et al 2015). We measured plant height in millimeters (mm) to the apical meristem of each plant using a ruler, and all floral traits were measured in mm with calipers after removing the corolla from each flower using forceps. We calculated herkogamy (stigma-anther separation) for each plant by subtracting the longest stamen length from the pistil length. Herkogamy acts as a metric of self-fertilization, where a smaller difference means a higher likelihood of self-fertilization (Karron et al. 1997).

### Statistical analyses

To understand whether the plastic and mean values of the same phenotype were genetically correlated, and therefore potentially subject to correlational selection, we performed a correlation analysis. We input all of our measured mean and plastic trait values into the R package *corrplot* which created a correlation matrix (Wei and Simko 2021). This allowed us to test, for example, the degree to which genotypes that expressed a highly lobed mean leaf shape in a long-day environment also displayed a large plastic change in leaf shape in response to different photoperiods.

To test for significant phenotypic plasticity and GxE in all *M. laciniatus* traits across photoperiods, we used a two-way ANOVA including genotype, photoperiod environment, and genotype by environment interaction as fixed effects (Lorts and Lasky 2020; Zhang and Lechowicz 1994). We calculated plasticity strength for each genotype by taking the difference between 15 and 11h mean phenotypic values of each genotype in each photoperiod (Valladares, Sanchez-Gomez, and Zavala 2006).

To examine patterns of trait variation consistent with local adaptation we tested for a relationship between mean trait expression and population elevation and latitude. We used an ANOVA with mean phenotype as the dependent variable for all traits measured. Elevation and latitude were fixed effects in our models while genotype was nested within population as a random effect. We used this same method to examine the relationship between the strength of trait plasticity and population elevation and latitude.

Reaction norm and environmental cline figures were created in R with the package *ggplot2* (Wickham 2016). Maps were created in R using the package *ggmap* as well as QGIS (Kahle and Wickham 2013; QGIS Development Team 2023).

## RESULTS

### Weak correlations between trait means and plasticity

We performed a correlation analysis to investigate whether the mean phenotype and plasticity of the same trait were genetically correlated and possibly subject to correlational selection. We found that at the individual genotype level mean trait expression was not tightly associated with the strength of plasticity in that same trait (Figure 3). Several mean traits were strongly correlated (*r* > +/-0.5) with one another. Leaf area, plant height and flowering time were all strongly correlated with each other as were corolla length, corolla width and stamen length. Mean leaf shape was most strongly correlated with mean leaf size (r = 0.59) and more weakly correlated with other mean traits. All floral plasticity measurements were tightly correlated with each other: stamen length (long and short), pistil length, corolla length and corolla width (*r* > +/-0.5). Due to the strong correlation between floral size traits we focus on corolla width as a representative floral trait for the rest of the results. For the majority of traits measured there was not a strong correlation between mean phenotype and plastic expression of the same phenotype (*r* < +/-0.5). For example, our results show that mean leaf shape expression of a genotype was only weakly correlated (*r* = 0.11) with the magnitude of leaf shape plasticity in that same genotype (Figure 3).

**Figure 3.**
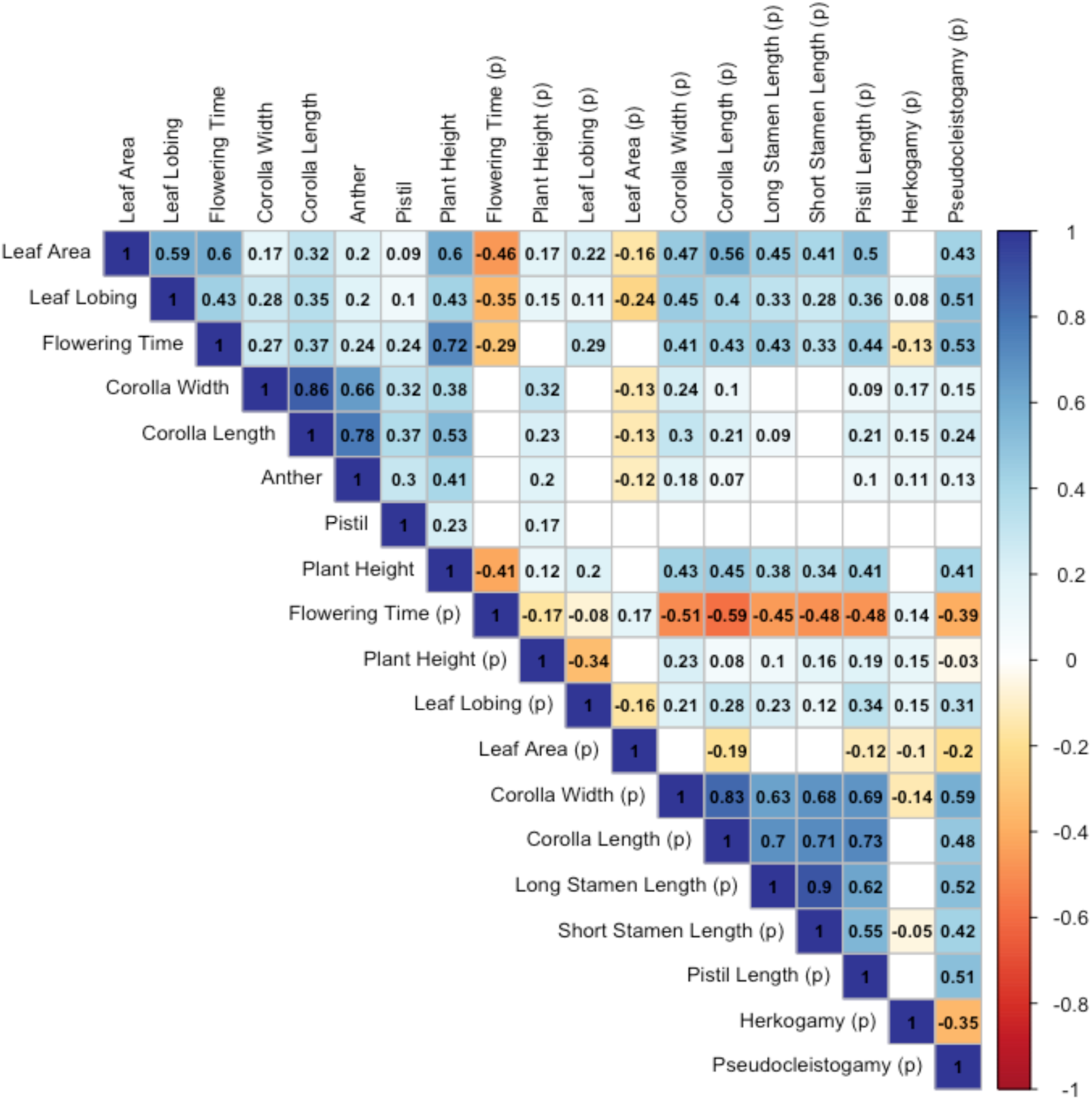
Correlation matrix of mean trait expression and plastic trait expression. Plastic trait expression is denoted by (p). Blank squares had an insignificant correlation (p > 0.05) when tested with a t-test.

### Genotypes from across the species’ range respond plastically to photoperiod

To test whether *M. laciniatus* exhibited plasticity in response to different photoperiods, we grew replicate genotypes in long and short day controlled environments. Significant phenotypic plasticity, measured by a significant effect of photoperiod environment in our models, was found in all traits that we studied (Figure 4, Table 2). Plant height showed the highest proportion of genotypes that exhibited significant plastic response to photoperiod change indicating that height in *M. laciniatus* is an environmentally labile trait. Across all genotypes the degree of plastic change in height was on average an increase in 45 mm between short and long days. Other phenotypes were less plastic at the species level in response to photoperiod such as leaf lobing, which on average only increased on our leaf lobing index by 0.03 between short and long days. However, a significant GxE interaction was detected in all traits studied indicating there is a great deal of genetic variation for phenotypic plasticity in this species (Table 2). In addition, some phenotypes had more similar patterns of plastic change than others, with over 40% of all genotypes growing taller, flowing faster, and with larger flowers in long days. This indicates that the direction of plasticity is mostly fixed across the geographic range of *M. laciniatus* in height, flowering time, and flower size. Genetic variation in leaf shape plasticity however was not as consistent in direction or degree across populations making it particularly interesting for follow up analyses. The detection of GxE in plant height, flowering time, floral size, and leaf shape makes these traits the focus of our remaining analyses testing for patterns of local adaptation within *M. laciniatus*.

**Figure 4.**
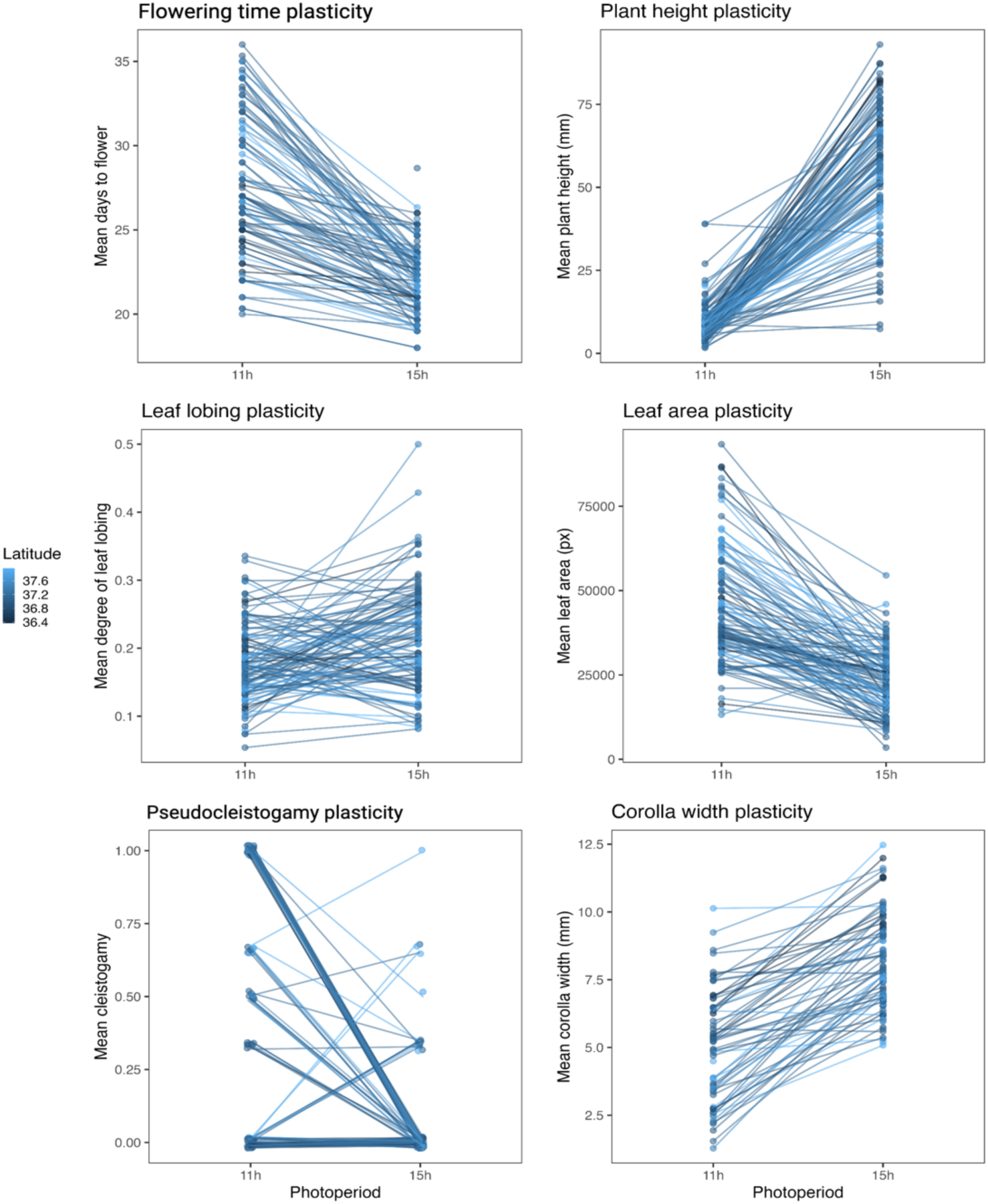
Reaction norms for a selection of focal traits. Trait values of *M. laciniatus* inbred lines were measured in controlled growth chamber conditions under two different photoperiods: 11 hour and 15 hour days. Reaction norms faceted by population can be found in Figure S2. See Table 2 for the significance of genotype by environment interactions for each trait.

**Table 2.** Significance of genotype (G), environment (E), and genotype by environment (GxE) interactions for each phenotype. A two-way ANOVA was used to generate these statistical comparisons: df = degrees of freedom, F = F-value, p = p-value.

| Trait | G | E | GxE |
| --- | --- | --- | --- |
| Plant height | df = 112<br>F = 4.0948<br>p < 0.001 | df = 1<br>F = 2251.7352<br>p < 0.001 | df = 109<br>F = 3.4516<br>p < 0.001 |
| Flowering time | df = 112<br>F = 8.47<br>p < 0.001 | df = 1<br>F = 809.2933<br>p < 0.001 | df = 88<br>F = 3.8501<br>p < 0.001 |
| Leaf lobing | df = 112<br>F = 3.3609<br>p < 0.001 | df = 1<br>F = 49.5744<br>p < 0.001 | df = 108<br>F = 2.0924<br>p < 0.001 |
| Leaf area | df = 112<br>F = 5.6028<br>p < 0.001 | df = 1<br>F = 600.5746<br>p < 0.001 | df = 108<br>F = 2.8207<br>p < 0.001 |
| Corolla length | df = 112<br>F = 5.7080<br>p < 0.001 | df = 1<br>F = 506.5164<br>p < 0.001 | df = 88<br>F = 2.8753<br>P < 0.001 |
| Corolla width | df = 112<br>F = 6.7879<br>p < 0.001 | df = 1<br>F = 699.9515<br>p < 0.001 | df = 88<br>F = 2.2367<br>p < 0.001 |
| Short stamen length | df = 112<br>F = 4.433<br>p < 0.001 | df = 1<br>F = 530.3004<br>p < 0.001 | df = 88<br>F = 2.8457<br>P < 0.001 |
| Long stamen length | df = 112<br>F = 4.9323<br>p < 0.001 | df = 1<br>F = 477.1790<br>p < 0.001 | df = 88<br>F = 2.4494<br>p < 0.001 |
| Pistil length | df = 112<br>F = 4.5903<br>p < 0.001 | df = 1<br>F = 409.9971<br>p < 0.001 | df = 88<br>F = 1.8443<br>p < 0.001 |
| Herkogamy | df = 112<br>F = 1.8216<br>p < 0.001 | df = 1<br>F = 104.4350<br>p < 0.001 | df = 88<br>F = 3.1339<br>p < 0.001 |
| Cleistogamy | df = 112<br>F = 2.7126<br>p < 0.001 | df = 1<br>F = 138.1892<br>p < 0.001 | df = 87<br>F = 3.5217<br>p < 0.001 |

### Mean leaf shape varies clinally with both elevation and latitude

To examine phenotypic patterns consistent with local adaptation in *M. laciniatus* we first tested for associations between variation in mean phenotype and elevation or latitude of origin. We found that mean phenotype expression was significantly associated with origin of population elevation in both morphological and life history traits of *M. laciniatus* (Table 3). Critical photoperiod, the photoperiod threshold required to reach flowering, was positively associated with population elevation of origin (R^2^ = 0.2, p = 0.003, Figure 5). Critical photoperiod is longer for high elevation populations, the majority of which did not flower in our 11 hour day treatment but did flower under 15 hour days. On the other hand, many genotypes from low elevation populations were able to flower under short days. Mean leaf shape also shows a significant positive association with population elevation, with leaves becoming more lobed at higher elevations (R^2^ = 0.36, p = 0.003, Figure 5). Two mean traits were marginally associated with population elevation in our first long day experiment: flowering time and corolla width (R^2^ = 0.11 and 0.12, p = 0.056 and 0.058, Figure 5). In these traits populations from higher elevations flowered later and had larger flowers than low elevation populations.

**Figure 5.**
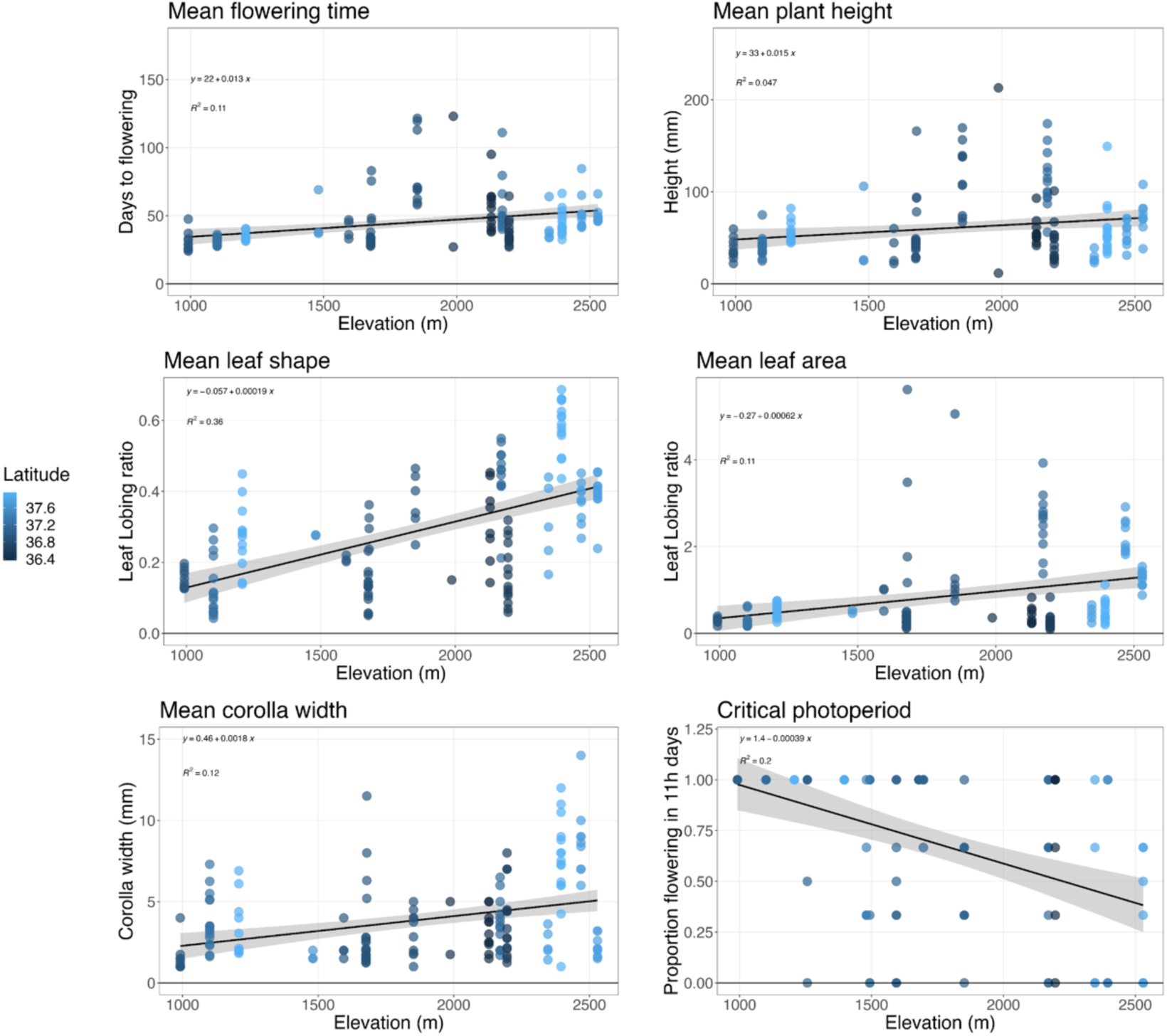
Trait mean expression per genotype plotted against elevation of origin for each population. See Table 3 for the statistical analysis of each clinal association.

**Table 3.** Significance of association between mean trait expression and population elevation. An ANOVA was used to generate these statistical comparisons. For all traits, numDF = 1, denDF = 18. *** = p < 0.01; ** = p < 0.05; * = p < 0.1.

| Trait | F-value | p-value | R<sup>2</sup> | Direction of correlation |
| --- | --- | --- | --- | --- |
| **Critical photoperiod** | **13.44429** | **0.003\*\*\*** | **0.2** | + |
| **Leaf shape** | **12.276596** | **0.003\*\*\*** | **0.36** | + |
| Flowering time | 4.179683 | 0.056* | 0.11 | - |
| Corolla width | 4.086909 | 0.058* | 0.12 | + |
| Pistil length | 2.57640 | 0.126 | 0.048 |  |
| Leaf area | 2.4211552 | 0.137 | 0.11 |  |
| Corolla length | 2.27840 | 0.149 | 0.094 |  |
| Herkogamy | 1.8451199 | 0.191 | 0.015 |  |
| Plant height | 0.397350 | 0.536 | 0.047 |  |
| Stamen length | 0.34039 | 0.5668 | 0.039 |  |

Mean leaf lobing was the only trait which had a relationship with latitude (R^2^ = 0.36, p = 0.0663; Figure 6). Even across the short latitudinal range of *M. laciniatus* we found that high latitude populations have more deeply lobed leaves than low latitudes. All other traits had no relationship with latitude (Figure 6, Table 4). Mean leaf lobing was therefore the only trait to be significantly associated with variation in both elevation and latitude across the species’ range.

**Figure 6.**
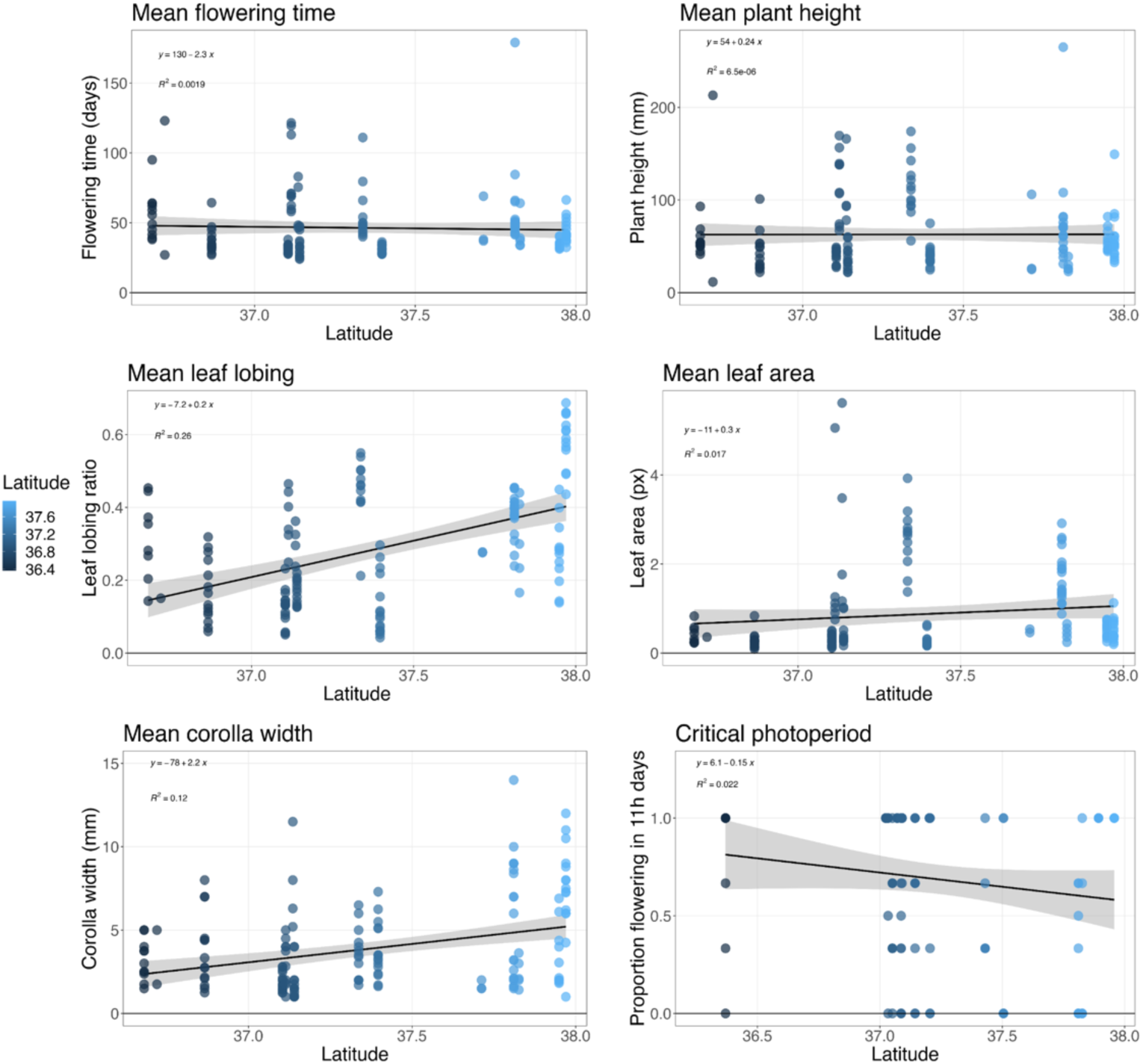
Trait mean expression per genotype plotted against latitude of origin for each population. See Table 4 for a statistical analysis of the clinal association of each trait with elevation.

**Table 4.** Significance of association between mean trait expression and population latitude. An ANOVA was used to generate these statistical comparisons. For all traits, numDF = 1, denDF = 13. *** = p < 0.01; ** = p < 0.05; * = p < 0.1.

| Trait | F-value | p-value | R<sup>2</sup> | Direction of correlation |
| --- | --- | --- | --- | --- |
| Leaf lobing | 4.017275 | 0.0663* | 0.26 | + |
| Corolla length | 1.837361 | 0.1983 | 0.15 |  |
| Flowering time | 1.753481 | 0.2083 | 0.0019 |  |
| Herkogamy | 1.535635 | 0.2372 | 0.00018 |  |
| Corolla width | 1.0265023 | 0.3295 | 0.12 |  |
| Anther length | 0.7934602 | 0.3892 | 0.056 |  |
| Critical photoperiod | 0.651162 | 0.4342 | 0.022 |  |
| Plant height | 0.6417213 | 0.4375 | 0.0000056 |  |
| Pistil length | 0.0470382 | 0.8317 | 0.016 |  |
| Leaf area | 0.0344051 | 0.8557 | 0.017 |  |

### Phenotypic plasticity in *M. laciniatus* varies only with elevation

We next set out to detect patterns consistent with local adaptation in phenotypic plasticity. Using a mixed model approach, we found that the strength of phenotypic plasticity in both life history and morphology was significantly associated with population elevation across the geographic range of *M. laciniatus* (Table 5). The degree of plastic change in leaf shape in response to our photoperiod treatments was greater in high than in low elevation populations (R^2^ = 0.26, p = 0.0663, Figure 7). High elevation populations became significantly more lobed in long days. Low elevation *M. laciniatus* populations had either very little change in leaf shape between treatments or the amount of lobing actually decreased slightly in response to long days. Therefore, both mean leaf lobing and leaf lobing plasticity increase in higher elevation populations. Plasticity in a number of floral size traits showed marginally significant trends (0.1 > p > 0.05) with elevation. The amount of phenotypic plasticity increased in higher elevation populations for long and short stamen length, pistil length, and pseudocleistogamy (Figure 7). Variation in flowering time and corolla size plasticity was not associated with elevation even though the mean values of these traits increased in higher elevation populations. Plant height, leaf size, and herkogamy plasticity also did not associate with altitude (Figure 7). When plasticity was analyzed for a relationship with latitude, no trait showed any clinal association (Figure S3, Table S2).

**Figure 7.**
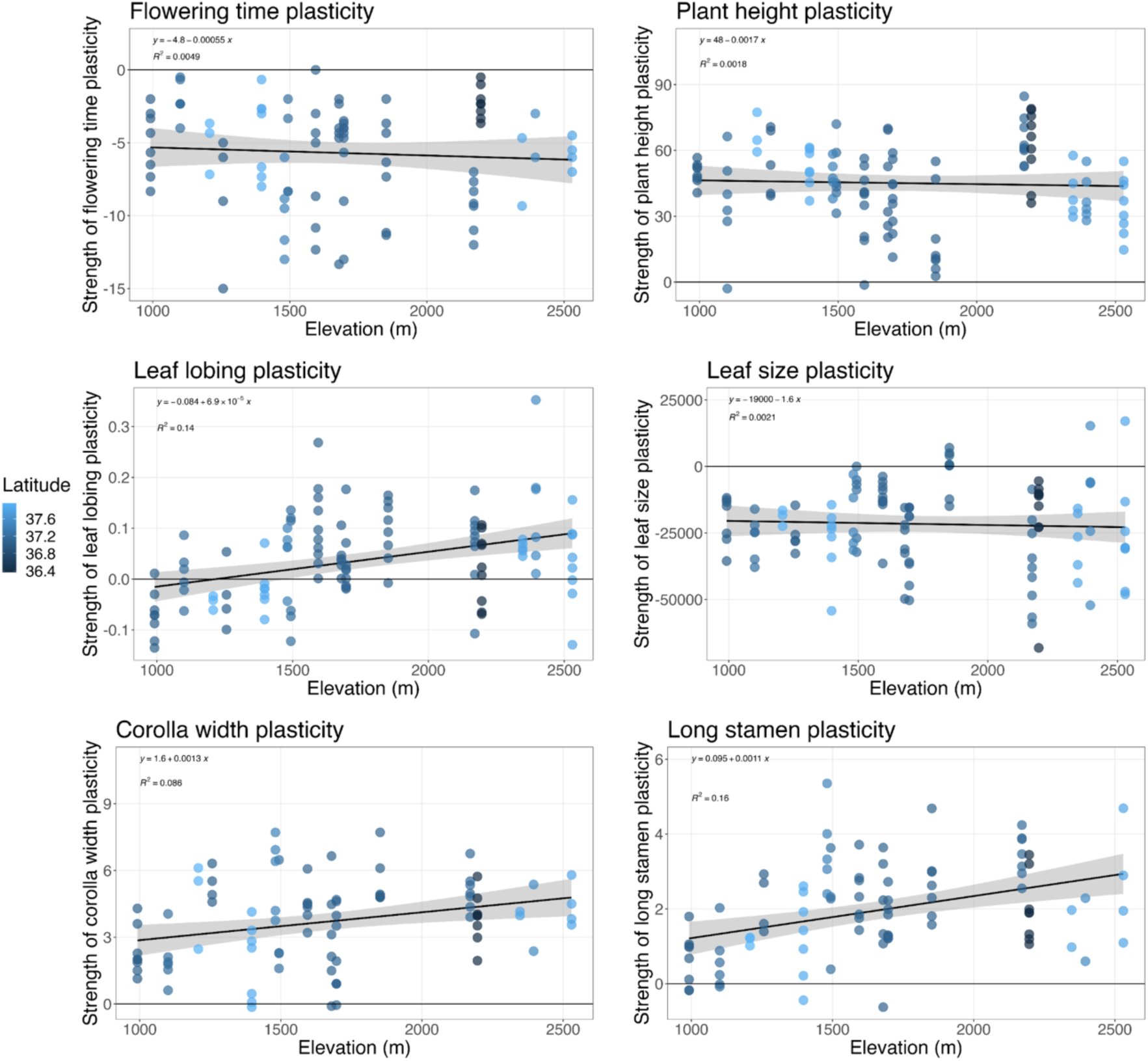
Mean trait plasticity per genotype plotted against elevation of origin for each population. See Table 5 for statistical analysis of the association between trait plasticity and latitude.

**Table 5.** Significance of association between strength of plasticity and population elevation. An ANOVA was used to generate these statistical comparisons. For all traits, numDF = 1, denDF = 14. *** = p < 0.01; ** = p < 0.05; * = p < 0.1.

| Trait | F-value | p-value | R<sup>2</sup> | Direction of correlation |
| --- | --- | --- | --- | --- |
| **Leaf shape plasticity** | **8.790705** | **0.010\*\*\*** | **0.14** | + |
| Short stamen length plasticity | 4.068938 | 0.063* | 0.14 | + |
| Pseudocleistogamy plasticity | 3.933972 | 0.0673* | 0.087 | + |
| Pistil length plasticity | 3.836726 | 0.070* | 0.13 | + |
| Long stamen length plasticity | 3.798262 | 0.072* | 0.16 | + |
| Corolla length plasticity | 3.7866308 | 0.117 | 0.1 |  |
| Corolla width plasticity | 1.746458 | 0.208 | 0.086 |  |
| Plant height plasticity | 0.365843 | 0.555 | 0.0018 |  |
| Herkogamy plasticity | 0.1446548 | 0.709 | 0.00000095 |  |
| Flowering time plasticity | 0.115048 | 0.740 | 0.0049 |  |
| Leaf area plasticity | 0.044490 | 0.836 | 0.0021 |  |

## DISCUSSION

In this study we investigated whether quantitative traits that are involved in adaptation to *M. laciniatus*’s harsh granite outcrop environment - flowering time, leaf shape and flower size (Ferris and Willis 2018) - also appear to be under selection in response to environmental variation across the species’ range. To test for patterns consistent with local adaptation to elevation and latitude we grew replicates of genotypes from across *M. laciniatus*’s geographic range under both long and short day lengths. Plasticity and GxE were detected in response to photoperiod in all traits that we measured. For local adaptation of plasticity to evolve there must be GxE in the traits’ expression and genetic variation in plasticity associated with variation in the environment. We found that patterns of genetic variation in the mean phenotypic expression of flowering time, flower size, and critical photoperiod were associated with elevation making them interesting traits to investigate in future studies about adaptation across *M. laciniatus’s* geographic range. In our study both mean and plastic leaf shape expression were significantly associated with elevation, strongly suggesting that these traits are involved in local adaptation within *M. laciniatus*. Furthermore, we found that leaf lobing expression in long days was positively associated with latitude as well as elevation indicating that increased lobing may be adaptive in colder environments.

### Mean phenotype and plastic trait expression seem genetically independent

In our study of range-wide phenotypic variation in *M. lacinaitus* mean trait expression had minimal influence on the degree of plasticity in the same trait (Figure 3). Notably in leaf lobing, which both mean and plastic variation are similarly associated with elevation there was little correlation between how lobed a leaf was in a long day environment and how plastic that genotype’s leaf shape expression was between day lengths. The independence of expression in these two traits implies that there is likely a separate genetic basis for mean and plastic expression of leaf lobing. Independence between mean and plastic trait expression has been theorized in a handful of studies (Ehrenreich and Pfennig 2016; Johansson et al. 2013), and identified in others (Lind et al. 2011; Snell-Rood et al. 2011; Tétard-Jones, Kertesz, and Preziosi 2011). While Snell-Rood et al. (2011) focus on an extreme form of plasticity, polyphenism, their study aimed to investigate their developmental divergence hypothesis first discussed in the literature with regards to plasticity by West-Eberhard (1989). Snell-Rood et al. (2011) found divergence in patterns of gene expression between different morphs in two species of horned beetle, *Onthophagus taurus* and *Onthophagus nigriventris*. Plasticity-biased genes were found to be more divergent than those with shared patterns of expression between beetle morphs. Tétard-Jones et al. (2011) also found independent plasticity QTL located separately from main effect QTL in barley growth, *Hordeum vulgare*, adding more evidence that plasticity is controlled by separate genetic loci than mean trait expression loci.

In a trait that is up- or down-regulated by common developmental pathways, expression of phenotypic plasticity is not linked to specific environmental cues and is more likely to be linked with mean trait expression (Snell-Rood et al. 2011). However, traits may also be controlled by environment-specific gene regulation, which adjusts expression in these common developmental pathways and can contribute to phenotypic plasticity. Given the lack of correlation between mean and plastic expression in *M. laciniatus* perhaps additional genes are triggered to cause differences in leaf shape in different environments. These adjustments would reduce pleiotropic constraints and in this way could facilitate rapid adaptation to different environments (West-Eberhard 1989, 2005).

### *Mimulus laciniatus* exhibits range-wide plasticity and GxE in response to photoperiod

Phenotypic plasticity is expected to evolve when future environmental conditions will vary, and presently sensed cues are reliable in predicting those future conditions (Snell-Rood and Ehlman 2021). We therefore may expect that plasticity evolved in response to *M. laciniatus*’ annual variation in climate and photoperiod, and was subsequently selected against in populations which have more stable, temporally homogeneous environments from year to year. We identified significant plasticity and GxE in all traits measured in different photoperiods in our set of range-wide *M. laciniatus* genotypes. Identifying GxE means there is genetic variation in plastic trait expression, and therefore selection can act on these different reaction norms potentially giving rise to local adaption.

Some traits such as plant height and flowering time appear to show a strong plastic response across genotypes, but less variation in strength and direction of plasticity among genotypes. Plant height and flowering time are strongly phenotypically correlated in our experiment (r = 0.72) which makes sense because these two phenotypes have been shown to be controlled by the same light sensing genetic pathway in other plants (Pigliucci and Schlichting 1996; Mockler et al. 2003). This is well documented in the *Arabidopsis* system, in which much is known about genetic controls linking flowering and plant height (Alvarez et al. 1992; Salehi et al. 2005). In *A. thaliana* the *terminal flower* and *FLOWERING LOCUS C* loci act antagonistically to genetic pathways that regulate a vegetative-to-reproductive switch in resource allocation. Given the likely genetic correlation this more universal plastic response in flowering and height across *M. laciniatus* populations may be due to genetic constraint (Auld, Agrawal, and Relyea 2010). Previous reciprocal transplant experiments have demonstrated that early flowering time is under strong selection in *M. laciniatus*’ harsh rocky outcrop environments (Tataru et al 2023; Ferris and Willis 2018). Given the ephemeral nature of water availability in the shallow rocky soils of this species’ habitat and the significant variation in the amount and timing of snow melt from year to year it seems likely that an ability to respond plastically in the timing of flowering is adaptive across *M. laciniatus’* range. Therefore, plasticity in crucial life history traits like flowering time and size could be universally, rather than locally, adaptive in our short-lived rocky outcrop specialist.

### Patterns of local adaptation to elevation in *M. laciniatus* present in both mean and plastic traits

Reciprocal transplant experiments have found selection for early flowering time, more highly lobed leaves, and larger flowers in *M. laciniatus’*s harsh granite outcrop habitat (Ferris and Willis 2018; Tataru, Wheeler, and Ferris 2023). However, it is not known whether these same traits are involved in local adaptation across *M. laciniatus’*s range. One environmental variable that is likely to drive population level adaptation in this system is variation in elevation across the Sierra Nevada, CA. Using a long day common garden we found that variation in critical photoperiod, mean flowering time, flower size, and leaf lobing as well as plasticity in leaf lobing was associated with population elevation. Previous studies have found low levels of gene flow between populations of *M. laciniatus* (Ferris, Sexton, and Willis 2014; Sexton et al. 2016), therefore associations between phenotypic variation and elevation that are repeated across multiple mountain clines may be more likely caused by natural selection than by gene flow alone.

In our sample of *M. laciniatus* populations critical photoperiod increases with elevation. This pattern could be indicative of local adaptation to flower under shorter day lengths that lower elevation populations experience during their early spring growing season. These results are supported by Friedman & Willis (2013) who previously found that flowering in higher elevation *M. laciniatus* populations requires a longer photoperiod. However, Leinonen et al. (2020) found that with a more limited population sample mid-elevation populations required the longest photoperiod to reach flowering. In addition, Leinonen et al. (2020) found no significant association of population elevation with other *M. laciniatus* phenological traits in response to photoperiod. These previous studies examined only a few populations of *M. laciniatus* (<6 populations) compared to our increased sampling of the species range (21 populations total). *Mimulus guttatus* is also known to have locally adapted to photoperiod, one such example being the geothermally adapted *M. guttatus* of Yellowstone National Park, WY. These plants have adapted to flower under short winter days when geothermal vents make the surrounding environment wet and warm, suitable for plant growth and reproduction (Lekberg et al. 2012). Photoperiodism has been documented in many plant species as a crucial cue for seasonal change (Davis 2002; Mockler et al. 2003; MacQueen et al. 2021). High elevation populations in other species have been found to be locally adapted to longer photoperiods similar to our findings, such as in *Populus* and red spruce tree species (Pauley and Perry 1954; Butnor et al. 2019) as well as more herbaceous alpine plant species such as *Arabidopsis arenosa* (Wos et al. 2022). High-elevation adaptation to longer photoperiod has also been identified in non-plant species, such as pitcher plant mosquitoes and burying beetles (Bradshaw and Lounibos 1977; Tsai et al. 2020). In turn, photoperiodism can inform how and when resources are directed during a plant’s life span (i.e. to growth or reproduction). In our study while short photoperiod leaves of *M. laciniatus* are less lobed, they are larger on average than long photoperiod leaves. For genotypes that didn’t flower in a short photoperiod, the resources were instead being shunted into production of vegetative tissue, which could be a resource conservation measure (Freschet et al. 2010; Reich 2014). A trait that allows for sensing time of year, such as photoperiodism would be beneficial for *M. laciniatus*, as it provides information to these plants on how much time there is left to grow in the season before the occurrence of seasonal drought and senescence in their ephemeral rocky environment.

Patterns of flowering time and corolla width are also consistent with local adaptation to elevation in *M. laciniatus* as higher elevation populations flower later and have wider corollas. The idea of later flowering being adaptive at high elevations is contradictory to what is typically expected in alpine and high latitude plants because of the shortened growing season in these environments. In most systems it is thought that populations at high elevation should flower faster than those at low because if individuals flower too late in the season they may be killed by an early frost before reproducing (Stock, Campitelli, and Stinchcombe 2014). However, *M. laciniatus*’s unique rocky outcrop environment combined with the Mediterranean climate of California produce a different expectation in this system that is tied to water availability rather than the timing of first frost. In the Sierra Nevada, CA the majority of seasonal soil moisture is due to snow melt with little rain occurring during the growing season. Snowpack levels are lower at low elevations meaning water resources will run out faster shortening the growing season due to drought at low vs high mountain sites. Early flowering time allows plants to escape from the onset of seasonal drought while bigger, later flowering plants usually occur in more continuously moist environments (Heschel and Riginos 2005; Hall and Willis 2006; Franks, Sim, and Weis 2007; Kenney et al. 2014; Ferris and Willis 2018). Therefore plants adapted to low elevations should flower faster in order to reproduce successfully compared to those at high elevations (Fox 1990). Dickman et al. (2019) and Leinonen et al. (2020) have also found that high elevation *M. laciniatus* flower later than low elevation populations. Flower size has been shown to increase in plant populations at high elevation due to the increased attractiveness of large flowers in an environment where pollinators are scarce (Fabbro and Körner 2004; Maad, Armbruster, and Fenster 2013). However, *M. laciniatus* is highly self-fertilizing so this explanation seems less relevant in our study system. Increased flowering time and larger flowers at high elevations where there is greater water availability may be adaptive in *M. laciniatus* because later flowering plants can become larger and more fecund. A large body of work exists in local adaptation of flowering time phenology and flower size, indicating that these traits often experience selection in nature in order to optimize fitness (Lowry and Willis 2010; Wang et al. 2014; Dickman et al. 2019; Leinonen et al. 2020).

Leaf shape, size, thickness, and number have all been studied previously in various plant species and identified as locally adaptive to harsh marginal habitats such as high light or extreme hot or cold temperatures (Nagy 1997; Dudley 1996; Parker, Rodriguez, and Loik 2003; Campitelli and Stinchcombe 2013; Ferris et al. 2015). In our study *Mimulus laciniatus* populations from higher elevations had more highly lobed and highly plastic leaves than populations from lower elevations (Figure 5, Figure 7). We also saw a positive trend between leaf lobing and latitude which suggests that it is adaptive to have more highly lobed leaves in northern populations. Lobed leaves have a thinner boundary layer which means they equilibrate more efficiently to air temperature through convection and therefore should not drop below ambient temperature on clear nights when leaves radiate heat to the sky (Givnish 1979). This could prevent lobed leaves from incurring frost damage when ambient temperature is near, but still above freezing (Schuepp 1993; Nobel 2009). Patterns consistent with increased cold tolerance of lobed leaves have been found in other plant systems. For example, along the east coast of North America there is a latitudinal cline in leaf shape in the Ivyleaf morning glory, *Ipomoea hederaceae*, with northern populations fixed for the lobed morph while those in the south have entire leaves (Campitelli and Stinchcombe 2013). When Campitelli, Gorton and colleagues (2013) tested for differences in leaf temperature they did find that lobed *Ipomoea* leaves stayed warmer at night than round leaves, although no significant difference in was found in freeze tolerance itself. Preliminary work on leaf temperature in *Mimulus* found that the lobed leaves of *M. laciniatus* stayed warmer throughout a 24-hour period than the round leaves of *M. guttatus* in common garden experiment in the field (K. Ferris unpublished data). A study using *Solanum lycopersicum* found that shade-avoidant plants had more complex leaf structure in both lobing and serration, indicating that high light can also drive a more complex leaf shape (Chitwood et al. 2015). The patterns we found of greater leaf lobing in both high elevation and high latitude populations of *M. laciniatus* are consistent with leaf lobing being adaptive in colder environments (Yapp 1912; Nobel 2009).

### Conclusion

Our study investigated how mean and plastic trait variation can contribute to local adaptation in heterogeneous environments. The evolution of leaf shape, critical photoperiod, and leaf shape plasticity across 21 *M. laciniatus* populations spanning multiple watersheds is consistent with a signature of local adaptation (James et al. 2023). The results of this study raise a number of questions for future research. First, are these patterns of mean and plastic trait expression truly due to natural selection? What genes underlie the plastic responses observed due to photoperiod? Are the genes that control for a plastic response in changing photoperiod the same that control mean trait expression in a long day environment, and does this hold true for all traits studied? What is the function of leaf lobing in the field setting? While functions are hypothesized above, there is currently a lack of data to validate these claims. In addition, by further studying the strength of selection on plastic traits it would be possible to investigate costs associated with plasticity in mismatched environments. Our data supports the idea that highly lobed leaf shape is locally adaptive in high elevation and northern populations of *M. laciniatus*. In adaptation between *Mimulus* species we see evidence that ephemeral water availability in *M. laciniatus’s* dry rocky habitat may be driving the evolution of lobed leaf shape (Tataru et al 2023; Ferris and Willis 2018). However high elevation highly lobed populations of *M. laciniatus* tend to have higher water availability than low elevation populations with less lobing, implying that the selective force causing local adaptation in leaf shape within this species may be different than the force driving divergent evolution between species. Future studies using genomic data in combination with experimental manipulation in the field will be able to further investigate these questions.

## ACKNOWLEDGEMENTS

We thank Natalie Gonzalez, Aditi Mahesh, and Cecilia Hammond for help with planting and data collection. We also thank Johanna Schmitt and Graham Coop for all of their support and feedback on the portion of this project completed at University of California Davis, and Mirielle Caton-Darby for assistance collecting the mean phenotypic data presented in this paper. This research was supported by the National Institute of General Medical Sciences of the National Institute of Health (NIH) under Award Number R35GM138224. The content is solely the responsibility of the authors and does not necessarily represent the official views of the NIH. This work was also supported by a Center for Population Biology Postdoctoral Fellowship at the University of California Davis and a Tulane University Ecology and Evolutionary Biology Graduate Student Grant.

For research conducted at UC Davis, we should take a moment to acknowledge the land on which we gathered. For thousands of years, this land has been the home of Patwin people. Today, there are three federally recognized Patwin tribes: Cachil DeHe Band of Wintun Indians of the Colusa Indian Community, Kletsel Dehe Wintun Nation, and Yocha Dehe Wintun Nation. The Patwin people have remained committed to the stewardship of this land over many centuries. It has been cherished and protected, as elders have instructed the young through generations. We are honored and grateful to be here today on their traditional lands.

For research conducted at Tulane University, we acknowledge and pay tribute to the original inhabitants of this land. The city of New Orleans is a continuation of an indigenous trade hub on the Mississippi River, known for thousands of years as Bulbancha. Native peoples have lived on this land since time immemorial, and the resilient voices of Native Americans remain an inseparable part of our local culture. With gratitude and honor, we acknowledge the indigenous nations that have lived and continue to thrive here.

## DATA AVAILABILITY

Data and code are available from the Dryad Digital Repository. https://datadryad.org/stash/share/xyNBULQagWnkmI3oAbWKPEIJcZPvxR-1xg3zxwGEnKI

## SUPPORTING INFORMATION

**Figure S1.**
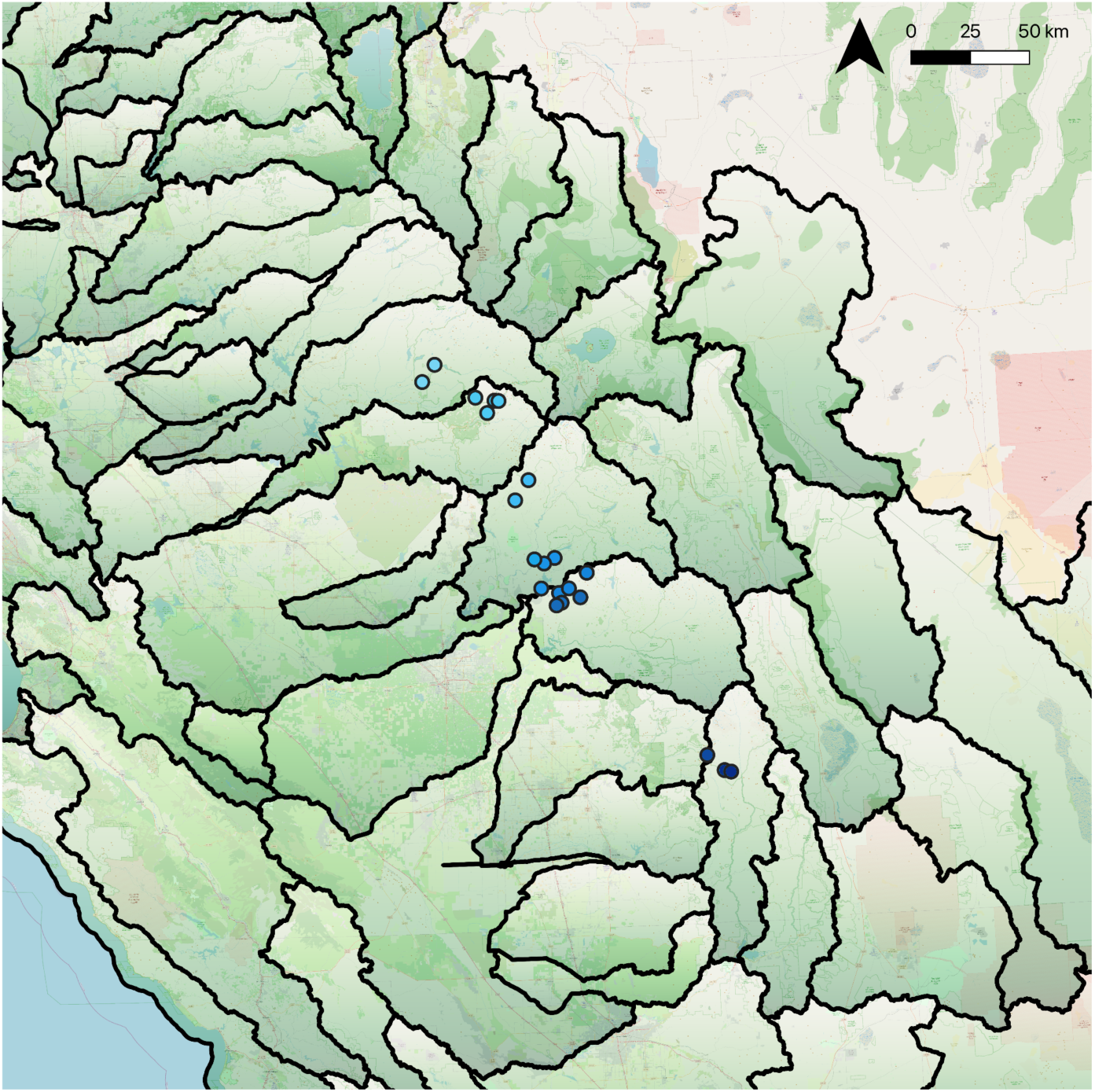
A map of study populations and watershed boundaries in California.

**Figure S2.**
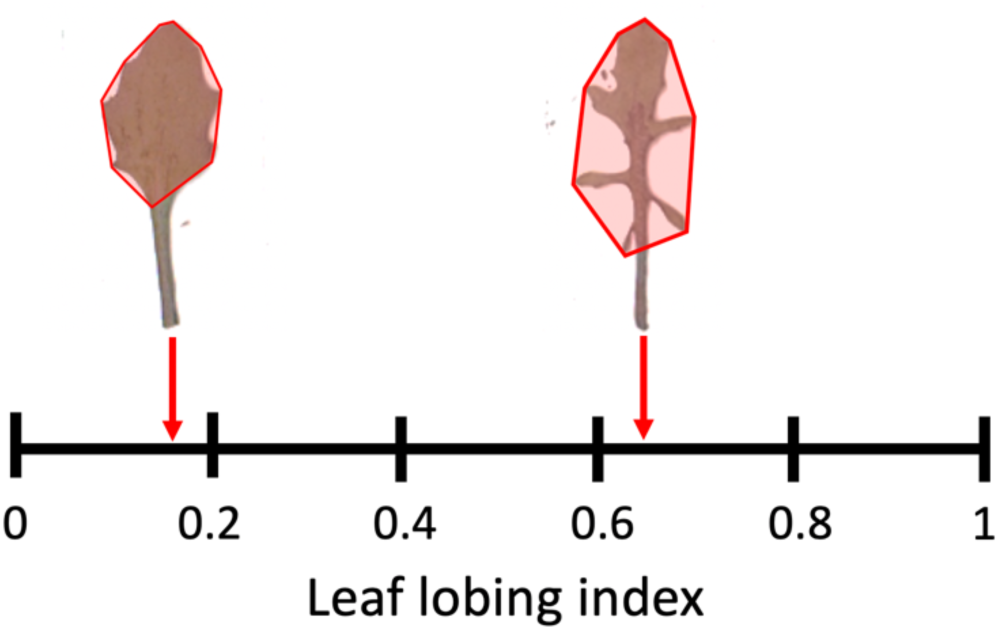
A visual depiction of the leaf lobing index used to measure leaf shape. The convex hull is outlined in red. A greater difference between true leaf area and the convex hull means a higher leaf lobing index. Leaf petioles were not included in leaf shape analysis.

**Figure S2.**
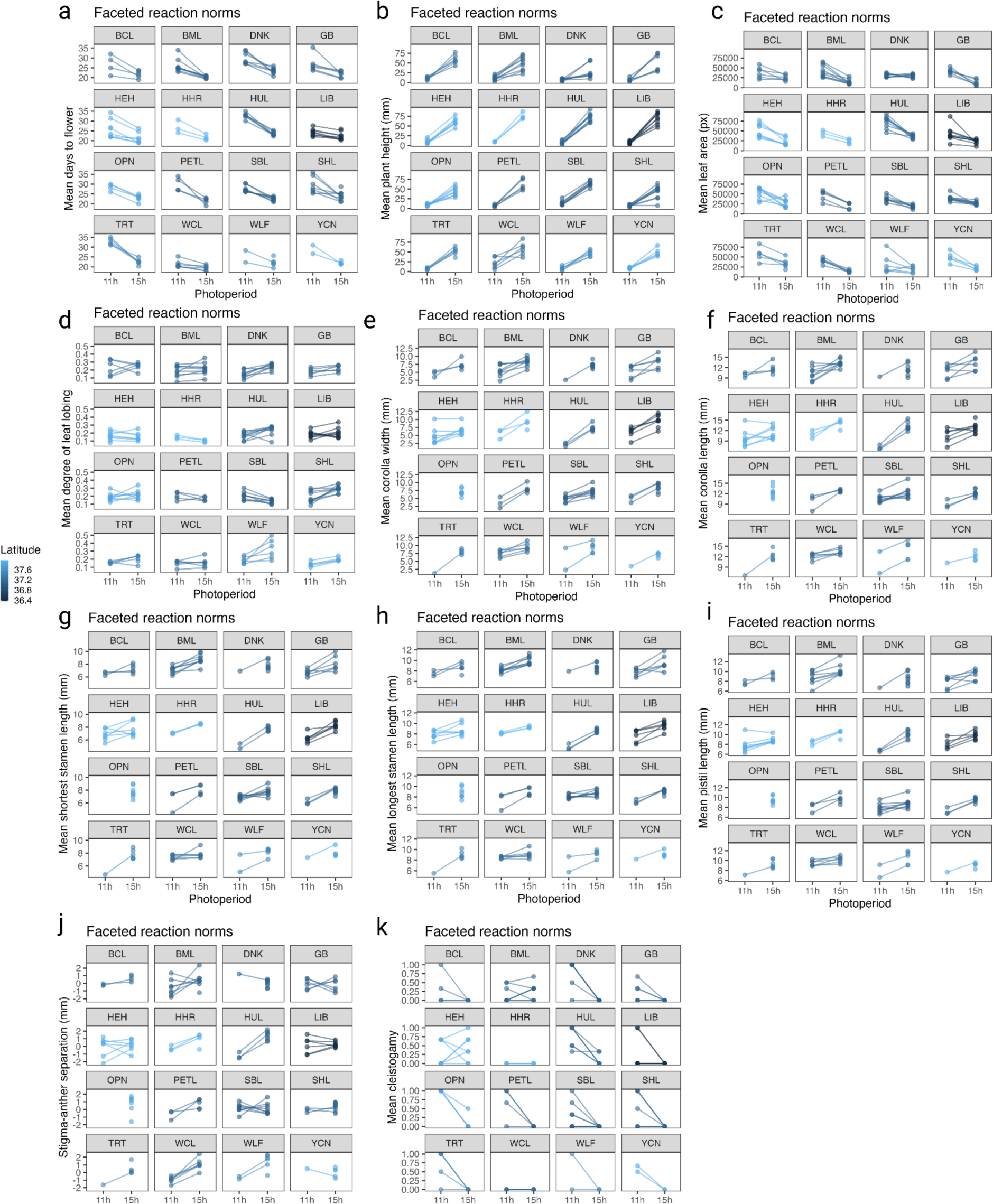
Reaction norms for each trait, faceted by population.

**Figure S3.**
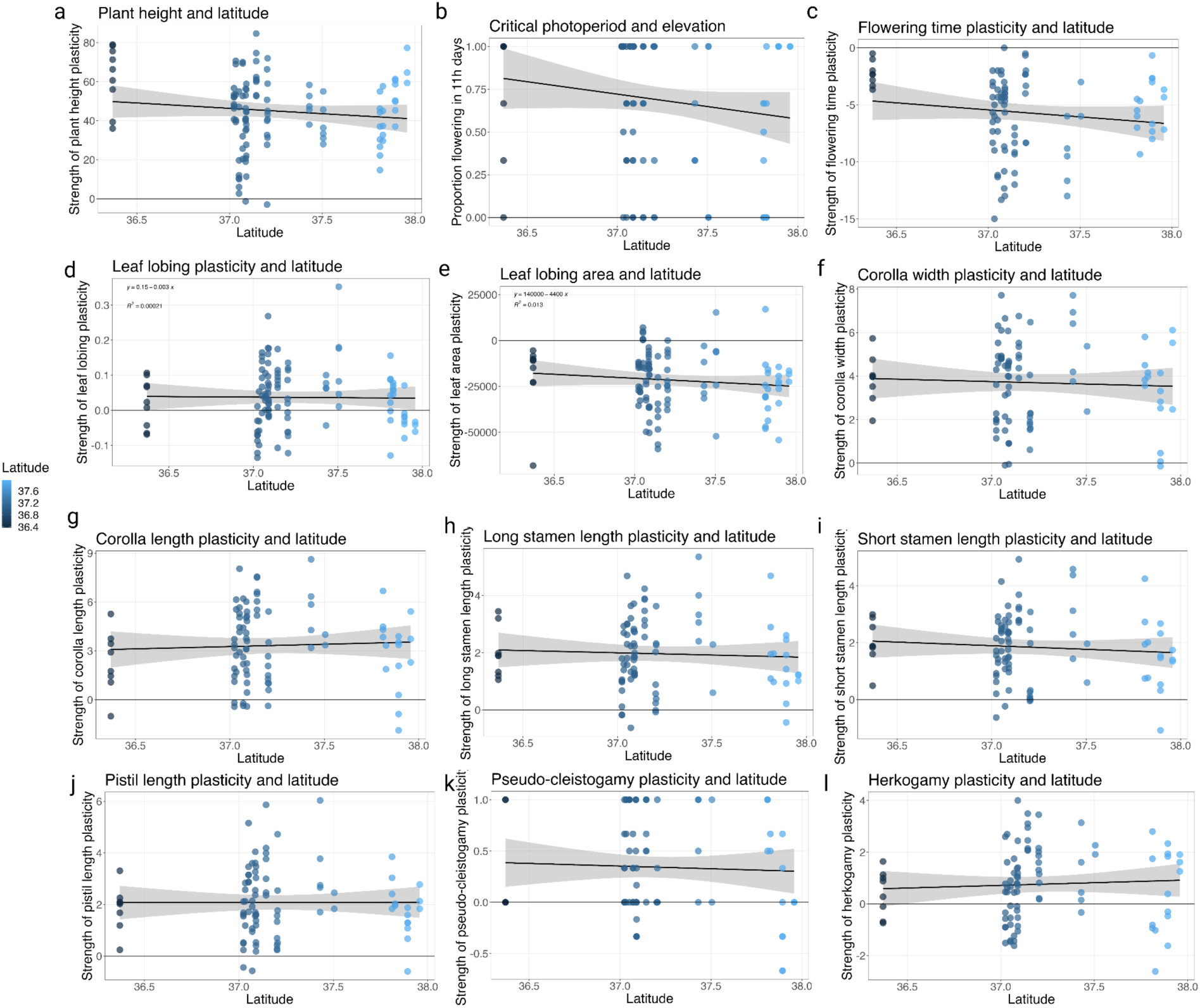
Trait plasticity mean expression per genotype plotted against latitude of origin for each population. See Table S2 for statistical analysis of clinal association.

**Table S1:** Each trait measured for mean and plastic phenotypes.

| Trait | Mean phenotype | Plastic |
| --- | --- | --- |
| Corolla length | Y | Y |
| Corolla width | Y | Y |
| Critical photoperiod | Y | -- |
| Flowering time | Y | Y |
| Herkogamy | Y | Y |
| Leaf area | Y | Y |
| Leaf lobing | Y | Y |
| Long stamen | Y | Y |
| Pistil length | Y | Y |
| Plant height | Y | Y |
| Pseudo-cleistogamy | -- | Y |
| Short stamen | -- | Y |

**Table S2.** Significance of association between plastic trait expression and population latitude. An ANOVA was used to generate these statistical comparisons. For all traits, numDF = 1, denDF = 13. *** = p < 0.01; ** = p < 0.05; * = p < 0.1.

| Trait | F-value | p-value | R<sup>2</sup> |
| --- | --- | --- | --- |
| Leaf area plasticity | 0.3967746 | 0.5397 | 0.013 |
| Flowering time plasticity | 0.3773531 | 0.5496 | 0.02 |
| Short stamen plasticity | 0.293991 | 0.5968 | 0.0072 |
| Long stamen plasticity | 0.217931 | 0.6483 | 0.0026 |
| Leaf lobing plasticity | 0.173843 | 0.6835 | 0.00021 |
| Corolla length plasticity | 0.0889135 | 0.7703 | 0.0025 |
| Plant height plasticity | 0.0513789 | 0.8242 | 0.014 |
| Pseudocleistogamy plasticity | 0.030234 | 0.8646 | 0.0019 |
| Pistil length plasticity | 0.00739 | 0.9328 | 0.0000064 |
| Herkogamy plasticity | 0.00497844 | 0.9448 | 0.0034 |
| Corolla width plasticity | 0.0027088 | 0.9593 | 0.0023 |

